# A coupled atrioventricular-aortic setup for in-vitro hemodynamic study of the systemic circulation: Design, Fabrication, and Physiological relevancy

**DOI:** 10.1101/2022.04.18.488661

**Authors:** Rashid Alavi, Arian Aghilinejad, Heng Wei, Soha Niroumandi, Seth Wieman, Niema M Pahlevan

## Abstract

In-vitro models of the systemic circulation have gained a lot of interest for fundamental understanding of cardiovascular dynamics and for applied hemodynamic research. In this study, we introduce a physiologically accurate in-vitro hydraulic setup that models the hemodynamics of the coupled atrioventricular-aortic system. This unique experimental simulator has three major components: 1) an arterial system consisting of a human-scale artificial aorta along with the main branches, 2) an artificial left ventricle (LV) sac connected to a programmable piston-in-cylinder pump for simulating cardiac contraction and relaxation, and 3) an artificial left atrium (LA). The setup is designed in such a way that the basal LV is directly connected to the aortic root via an aortic valve, and to the LA via an artificial mitral valve. As a result, two-way hemodynamic couplings can be achieved for studying the effects that the LV, aorta, and LA have on each other. The collected pressure and flow measurements from this setup demonstrate a remarkable correspondence to clinical hemodynamics. We also investigate the physiological relevancies of isolated effects on cardiovascular hemodynamics of various major global parameters found in the circulatory system, including LV contractility, LV preload, heart rate, aortic compliance, and peripheral resistance. Subsequent control over such parameters ultimately captures physiological hemodynamic effects of LV systolic dysfunction, preload (cardiac) diseases, and afterload (arterial) diseases. The detailed design and fabrication of the proposed setup is also provided.

## 1. Introduction

The performance of the human circulatory system cannot be fully comprehended without understanding the interactions of the left ventricle (LV), the left atrium (LA), and the arterial system [1]. The abnormal coupling of the LV-aorta system and the LV-LA system has been shown to contribute to the pathophysiology of system-level cardiovascular diseases [2–6]. In healthy conditions, there exists an optimal hemodynamic balance between the LV, the arterial network (aorta and its vascular branches), and the LA that guarantees delivery of the cardiac output with minimal energy expenditure and modest pulsatility in flow and pressure [2, 6–10]. Optimal atrioventricular-aortic hemodynamic coupling can be impaired due to age-related or disease-related changes. For example, recent clinical studies have shown that stiffening of the proximal aorta is associated with pulsatile load on the heart and can lead to the development of heart failure [3, 6, 11, 12]. While the physiological importance of such optimal hemodynamic balance between the LV, the LA, and the vascular network has been shown extensively [2, 3, 7, 13–15], the effects of the individual contributions from ventricular, atrial, and arterial parameters towards generating pressure and flow waves remain unexplored. In order to study the isolated effects of such parameters, it is essential to keep other parameters of the circulation either constant or under control.

In-vitro fluid dynamics studies of cardiovascular systems have shown to be effective in understanding the underlying hemodynamics mechanisms of diseases [4, 16–21]. These setups can reduce the high expenses and risks associated with clinical trials, can facilitate the test of diagnostic and therapeutic devices, and can be used to validate computational models of physiological phenomena [22–24]. In addition, such models allow one to study one parameter at a time while controlling all other parameters in a physical setting. This provides a unique advantage over pre-clinical models (e.g., rat models), where complex physiological interactions and biological variabilities exist [17]. Previous studies have introduced various in-vitro hemodynamic simulators and have demonstrated the physiological relevancies of their setups [4, 17, 26, 27]. However, none of these in-vitro simulators account for direct two-way coupling between both the LV-aorta and LV-LA subsystems.

This manuscript introduces a novel in-vitro hydro-mechanical system that is well-suited for studying hemodynamic couplings of the LV-arterial system. This experimental setup is designed to investigate the pulsatile hemodynamics and arterial wave reflections in the (human-scale) systemic circulation. In the setup, the main determinants of cardiovascular function can be controlled by: *i*) the contractile state of the LV; *ii*) the afterload (determined by vascular stiffness and the total peripheral resistance (TPR)); and *iii*) the LV preload (quantified by left ventricular end-diastolic pressure (LVEDP)). In our design, the basal LV is directly connected to the aortic root on the arterial side and to the artificial left atrium (LA) on the atrial side, with aortic and mitral valves in between. In this manuscript, the design and fabrication of the proposed setup are described in detail. The physiological accuracy and relevancy of the experimental setup are also discussed across a wide range of healthy and diseased conditions.

## 2. Materials and method

### 2.1. Description of in-vitro experimental setup

#### Hydraulic Circuit Components

The main components of our in-vitro experimental setup are *i*) the atrioventricular simulator; *ii*) the aortic simulator; *iii*) the total arterial resistance and compliance simulator; and *iv*) the venous simulator. The atrioventricular simulator consists of a compliant LV sac that is connected to the aortic root on one side via an artificial aortic valve and that is connected to the artificial LA on the other side via an artificial mitral valve (*Medtronic MOSAIC® 305 CINCH®*). The LV sac employed in this simulator is installed inside an LV chamber that connects the artificial LV to a computer-controlled piston-in-cylinder pulsatile pump (*ViVitro* Labs Inc, SuperPump, AR SERIES). The aortic simulator includes the artificial aorta and the end-organ simulators at the terminus of each branch, the latter of which consist of half-filled air syringes and clamps mounted at the outlets in order to account for the compliance and resistance of the eliminated vasculatures. The total arterial resistance and compliance simulator consists of a pair of half-filled air chambers from acrylic glass connected to each other with a resistance valve in between. In order to control the compliance of the system through the compliance chambers, pairs consisting of a pressure bulb (for applying the pressure) and a pressure gauge (for monitoring of the pressure) are mounted on top of each chamber. The height of the air column inside the compliance chambers, as well as the syringes, are adjusted as has been described in previous work [4]. The venous reservior consists of an open tank that is connected to the LA.

#### Hydraulic Circuit Functioning

The compliant LV sac is contracted (squeezed) inside a plexiglass container using a pulsatile pump programmed by the ViVigen interface (*ViVitro* Labs Inc.). Using physiological input waveforms for the piston pump displacement (as shown in S1 File), the fluid outside of the LV sac is pressurized (LV contraction) and depressurized (LV relaxation) as the piston moves. After the pressure starts to rise inside the LV sac, the prosthetic aortic valve opens, and the test fluid enters the aorta from the LV and begins to generate hemodynamic waves that propagate down the aorta. The fluid is then exited from the aorta and its branches via the two compliance chambers that are installed at the end of the aortic loop. The testing fluid then flows from the two compliance chambers into the reservoir tank, which then flows towards the artificial LA. As the LV starts to relax (due to the depressurization inside the LV chamber), the mechanical mitral valve opens, and the fluid subsequently pumps back inside the LV. This completes the in-vitro cardiac loop. A schematic of the full hydralutic circuit corresponding to the left atrioventricular-aortic hemodynamic simulator setup and its corresponding circulation path is presented in Fig. 1.

**Figure 1.**
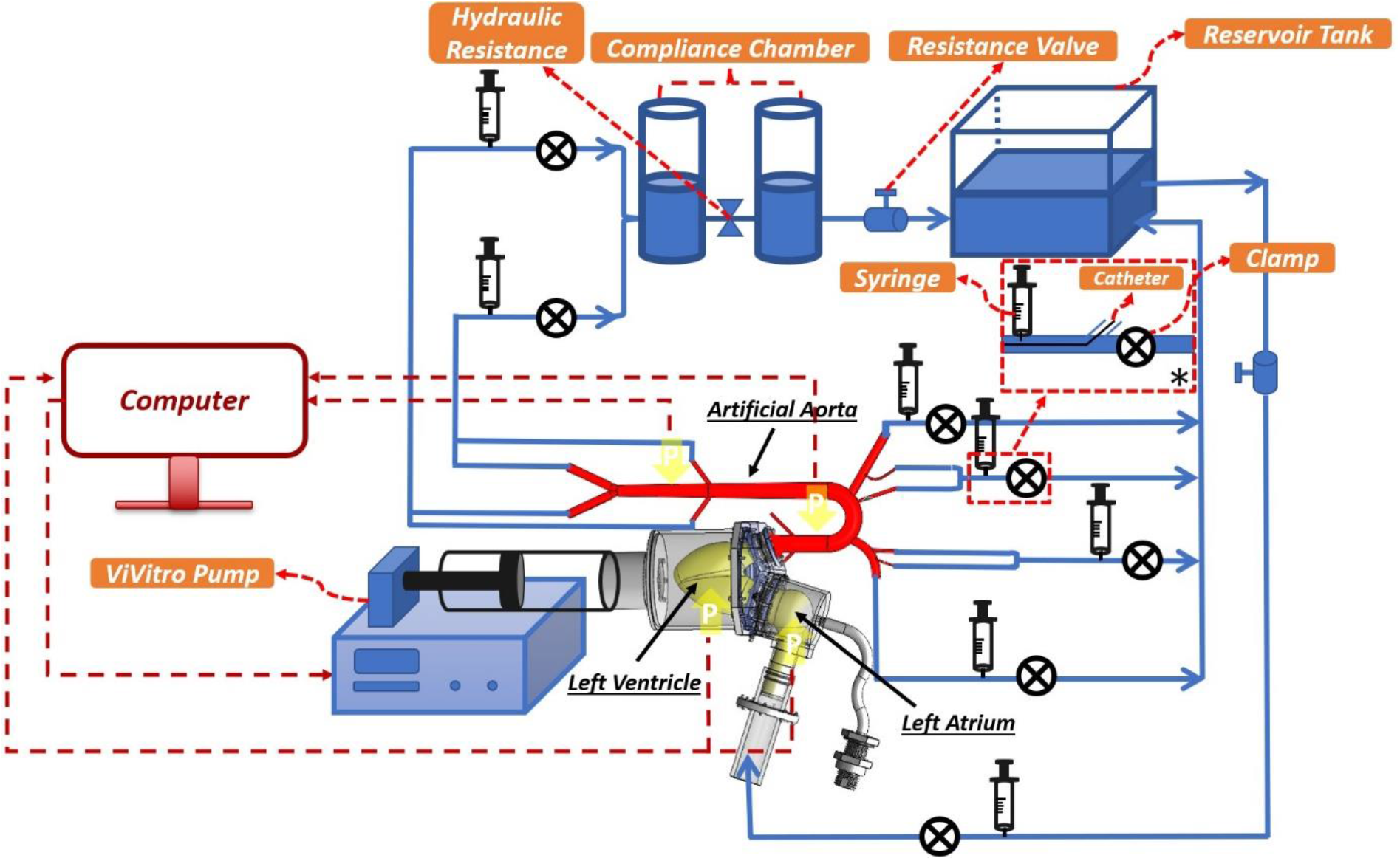
Schematic of the full hydralutic circuit corresponding to the left atrioventricular-aortic hemodynamic simulator setup and its circulation path. The schematic includes the complete in-vitro circulation system consisting of the LV, the aorta, the vascular components of the systemic circulation, and the left atrium (*: schematic of a sample outlet unit).

#### Design considerations for LV-aorta and LV-LA hemodynamic couplings

In the design process, we have ensured that the two-way couplings between the LV, LA, and aorta are captured by the in-vitro setup. As such, for the LV-aorta interface design, the basal LV is directly coupled with the aortic root through a prosthetic aortic valve. The LV-aortic coupling is machined from rigid acetal plastic that also serves as housing for the aortic valve (Fig. 2a). In the LV-LA interface design, the artificial LV is coupled with the artificial LA on the atrial side through a mechanical mitral valve. The LV-LA coupling is also fabricated from rigid acetal plastic and serves as housing for the mitral valve (Fig. 2a). Each of the coupling pieces mount to a bulkhead on the LV-housing with an O-ring sealed flange. Grooved sleeves extending from either side of the coupling flanges provide sealing surfaces for the compliant cuffs of the LV, LA, and aorta. O-rings that are placed under tension around the cuffs at the grooved portion of the sleeves complete this seal. The mechanical or prosthetic valves are sealed into the couplings by compressing their suture rings against inner rims on the couplings with a threaded retaining ring. The LV-aorta coupling includes a Luer-lock compatible port (Fig. 2b) through which a catheter can be inserted for measuring LV internal pressure. Efforts were made to keep the couplings as short as possible in order to minimize usage of rigid material along the LA-LV-aorta flow path. This coupling/valve housing approach is easily adapted to different LV, LA, and aorta sizes as well as a variety of mechanical or prosthetic valve types. Overall, the two-way couplings are mainly provided via the artificial valves and direct atrioventricular-aortic couplings with minimal inter-connectors and a cohesive media.

**Figure 2.**
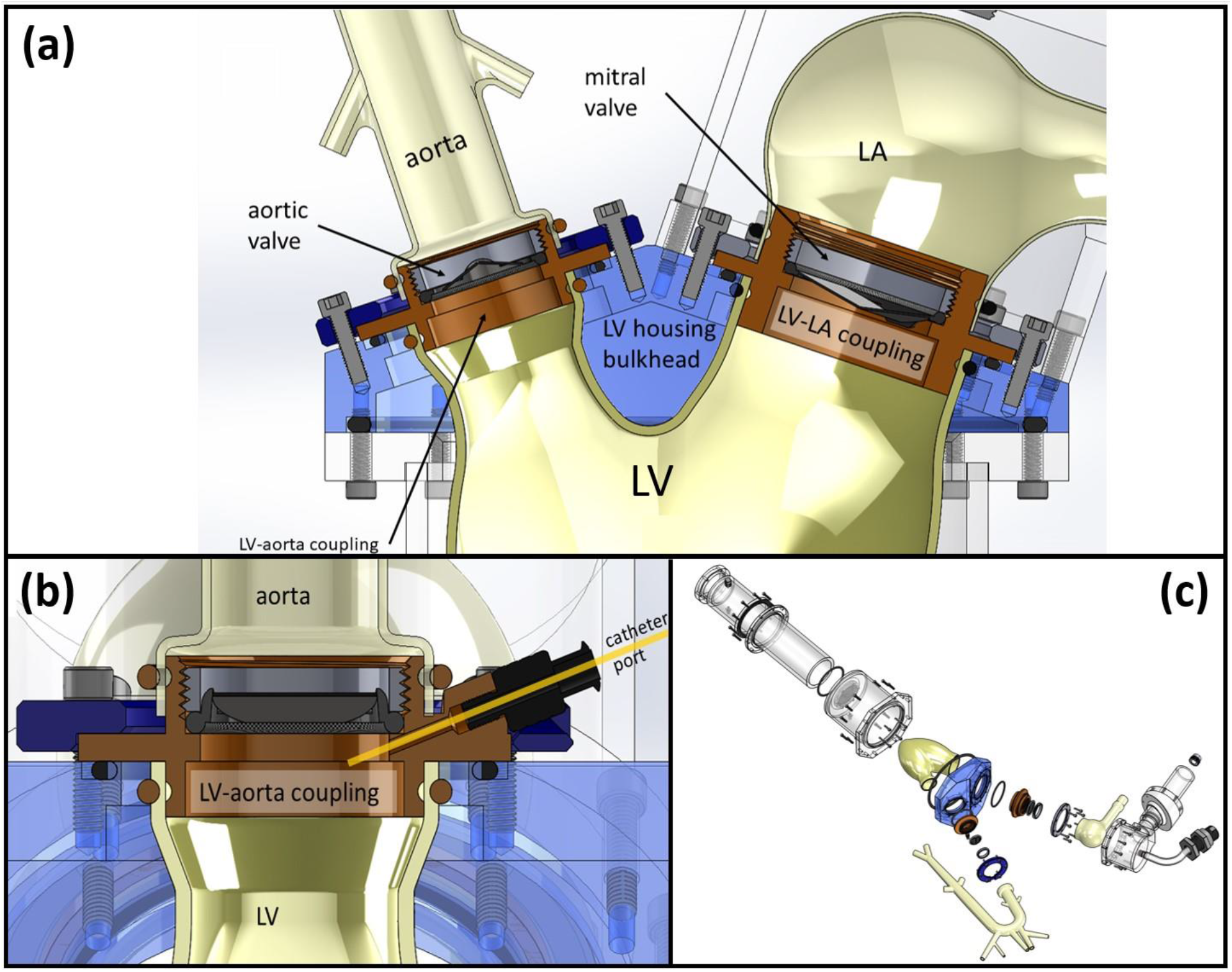
Detailed schematics of the LV-LA and the LV-aorta couplings. **(a)** Acetal plastic inserts placed in the LV housing bulkhead serve as both LV-LA and LV-aorta couplings as well as housings for their respective mitral and aortic valves. **(b)** A cross-sectional view of the LV-aorta coupling illustrates how the LV-aorta coupling insert includes a duct through which a catheter may be inserted for LV internal pressure measurements. **(c)** Exploded view of the setup components corresponding to the LV-aorta and LV-LA coupling mechanisms.

A picture of the final fabricated setup is shown in Fig. 3. The detailed CAD design of the constructed setup (3D CAD files) can be found in the manuscript’s Supplementary Material (see S2 File).

**Figure 3.**
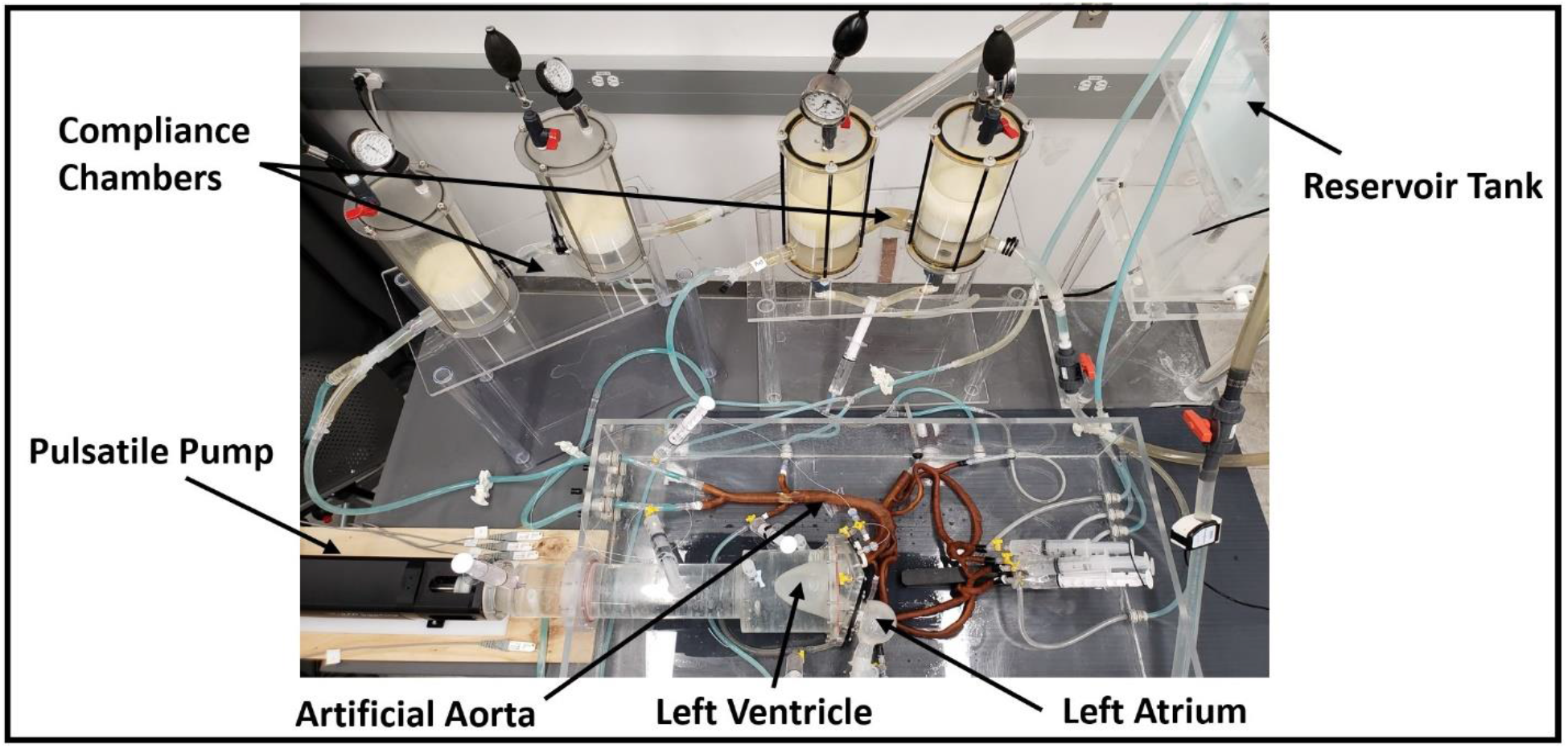
Picture of the complete in-vitro circulation system consisting of the LV, the aorta, the vascular components of the systemic circulation, and the left atrium.

### 2.2. Artificial organ fabrication

The artificial organs installed in the setup (e.g., aorta and LA) are fabricated using natural latex rubber (*Chemionics Corp.*) and silicone rubber (RTV-3040, *Freeman Manufacturing & Supply Company*). These materials are chosen based on their characteristic ability to replicate the stiffness of a physiological aorta [4]. The artificial aortas are fabricated in-house based on the one-to-one human-scale molds consisting of the ascending aorta, the aortic arch, the thoracic aorta, the abdominal aorta, and all the major branches (e.g., coronary and renal arteries). The aortic molds are built using either polyvinyl alcohol (PVA) via a 3D printer (*Ultimaker S5 Dual Extrusion*) or using a stainless-steel metal mold. Figure 4 and Table 1 list the segments and corresponding dimensions of the aortic molds that are employed in this work.

**Figure 4.**
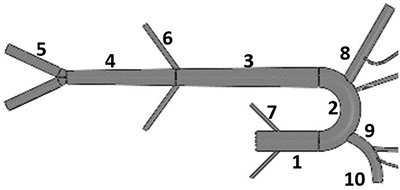
Schematic and segment diagram of the aorta mold.

**Table 1.**
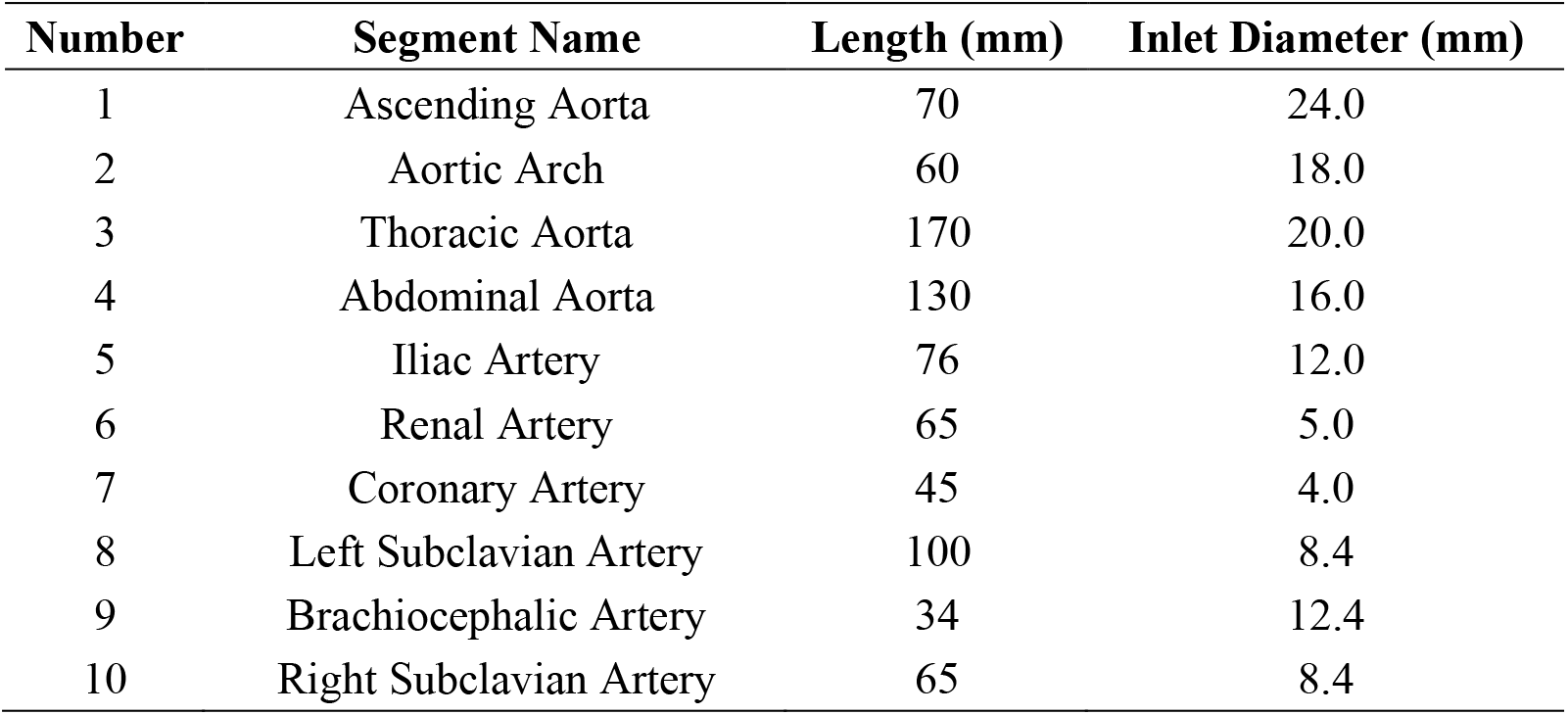
Geometric characteristics of the aorta mold.

| Number | Segment Name | Length (mm) | Inlet Diameter (mm) |
| --- | --- | --- | --- |
| 1 | Ascending Aorta | 70 | 24.0 |
| 2 | Aortic Arch | 60 | 18.0 |
| 3 | Thoracic Aorta | 170 | 20.0 |
| 4 | Abdominal Aorta | 130 | 16.0 |
| 5 | Iliac Artery | 76 | 12.0 |
| 6 | Renal Artery | 65 | 5.0 |
| 7 | Coronary Artery | 45 | 4.0 |
| 8 | Left Subclavian Artery | 100 | 8.4 |
| 9 | Brachiocephalic Artery | 34 | 12.4 |
| 10 | Right Subclavian Artery | 65 | 8.4 |

Dipping and coating casting methods are used for the natural latex and silicone-based fabrications, respectively. For the latex-based aortas, the following fabrication steps are taken: *i*) dipping the aortic mold inside the liquid latex container; *ii*) removing the coated mold from the container; *iii*) curing the coated material for 2 hours in standard room temperature (25°C); *iv*) repeating the coating for more layers (as necessary) so as to achieve the desired aortic compliance (steps *i* to *iii*). For the silicon-based fabrications, the following steps are taken: *i*) mixing the catalyst and the base of the silicon rubber (RTV-3040, *Freeman Manufacturing & Supply Company*) with a mass ratio of 10 (base) to 1 (catalyst); *ii*) removing the mixture bubbles using a vacuum pump chamber; *iii*) coating a light silicon sheet using 25 g of the mixed solution using a soft-tip acrylic brush; *iv*) curing the coated material for 16 hours at standard room temperature (25C); *v*) repeating the coating for more layers (as necessary) so as to achieve the desired aortic compliance (steps *i* to *iv*). For ensuring surface uniformity, the mold is turned upside down at each drying step. Table 2 presents the measured pulse wave velocities (PWV) and aortic compliances (AC) of the final fabricated aortas employing either silicone or latex. AC is measured by adding incremental volumes of fluid and measuring the corresponding incremental change in pressure. Details on acquiring PWV measurements are described in Section 2.3.

**Table 2.** Dynamic and physiological properties of the fabricated aortas.

| Aorta No. | PWV (m/s) | AC (mL/mmHg) | Material |
| --- | --- | --- | --- |
| Aorta-1 | 5.6 | 2.28 | Silicone |
| Aorta-2 | 10.6 | 1.78 | Latex |
| Aorta-3 | 12.6 | 1.51 | Latex |
| Aorta-4 | 16.6 | 1.43 | Latex |
| Aorta-5 | 17.3 | 1.31 | Latex |
| Aorta-6 | 18.0 | 1.19 | Latex |
| Aorta-7 | 19.7 | 0.83 | Latex |
| Aorta-8 | 21.7 | 0.71 | Silicone |

After casting the models using either latex or silicone, the prepared aortas are removed from their respective molds. For the metal molds, removal is achieved through gentle injection of water at the material/mold interface. For PVA molds, removal is achieved by immersing the models inside water at standard room temperature for 96 hours. The PVA mold is then dissolved in the water, and the PVA residue is manually detached from the latex or silicon aorta at the end of the process. Lastly, to enhance surface quality, the latex aortas are submerged in Clorox bleach for 12 hours, followed by another 12 hours of submerging in water for the final curing process. The same procedures are used for fabrication of other artificial organs (e.g., LA) with the silicone-based process described in detail earlier in this section, employing one-to-one human-scale PVA molds.

### 2.3. Procedures and Measurements

The in-vitro atrioventricular-aortic simulator has been tested across a wide range of physiological hemodynamic conditions (e.g., heart rate (HR)=50 to 125 bpm, cardiac output (CO)=2 to 5 L/min, LVEDP ≈ 0 to 25 mmHg, aortic systolic blood pressure (SBP) ≈ 80 to 170 mmHg, aortic diastolic blood pressure (DBP) ≈ 55 to 90 mmHg) [36–39]. In order to account for the changes in LVEDP, the venous return pressure is increased by adjusting the height of the venous reservoir tank connected to the LA. For assigning cardiac parameters to the setup, user-defined input waveforms are imported into the pulsatile pump controller (ViViGen interface). The frequency of the operation for the pump (which determines the HR) is also modified using the Vivigen interface on the computer unit. The input profile of the pulsatile pump system is adjusted to simulate the impact of LV contractility on system hemodynamics. The simulated contractility ranges from LV-*dp/dt_Max_*=937.4 mmHg/s to *LV-dp/dt_Max_*=2558.3 mmHg/s, covering low to high contractility conditions. Using resistance clamps connected to the end-organ outlets, the cross-sectional area of the outlet units is controlled to change the TPR of our atrioventricular-aortic simulator. In order to evaluate the physiological relevancy of the atrioventricular-aortic setup under normal condition, we simulate a baseline of *HR* = 75 bpm, *CO* = 5 L/min, LVEDP = 7 mmHg, SBP = 121.20 mmHg, DBP = 80.78 mmHg, AC = 1.51 mL/mmHg, and *TPR* = 18.85 mmHg · min/L.

For modeling LV systolic dysfunction, we run the setup under reduced CO conditions. This is achieved by reducing the stroke volume of the LV through adjusting the displacement amplitude of the LV-excitation pump. In order to simulate age-related or disease-related alterations in aortic stiffness, different physiological stiffnesses are achieved by changing the number of applications of dipping (for latex aortas) or coating (for silicone aortas). The corresponding aortic stiffness is quantified by measuring the PWV (the speed at which hemodynamic waves propagate inside the vasculature). Physiologically speaking, PWV depends on the structural alterations (which can be induced by aging) and transient functional changes in the arterial wall [28]. To compute the PWV shown in Table 2, the foot-to-foot method is applied for each one of the aortas using the measured pressures inside the setup. In such a method, the delay time is measured from the foot of the first propagating wave at the aortic root to the foot of the second propagating wave at the aortic bifurcation. The distance between the two sites is a known quantity that is based on the corresponding design of the aortic models.

A list of the parameters that are measured in this study, as well as the devices that have been used for the process of measurement, include: *i*) pressure (via a Millar MIKRO-TIP® Catheter Transducer (*Mikro-Cath*) using a PowerLab 4/35 from ADInstruments); *ii*) flow (via a Transonic Flowmeter (TS410)); and *iii*) heart sound (via a non-invasive wireless optical tonometer called Vivio [29]). In order to assess the effects of each input variable (e.g., LV contractility) on the local and global cardiovascular performance, four measurement sites are selected for the hemodynamic data collection: the central LV, the ascending aorta (6 cm away from the aortic root), the abdominal aorta (23 cm away from the aortic arch center), and the central LA. Water is used as the circulating fluid in all experiments, and any visible air bubbles are removed prior to running experiments. Table 3 lists the range of parameters employed to simulate realistic hemodynamic conditions in our in-vitro simulations.

**Table 3.**
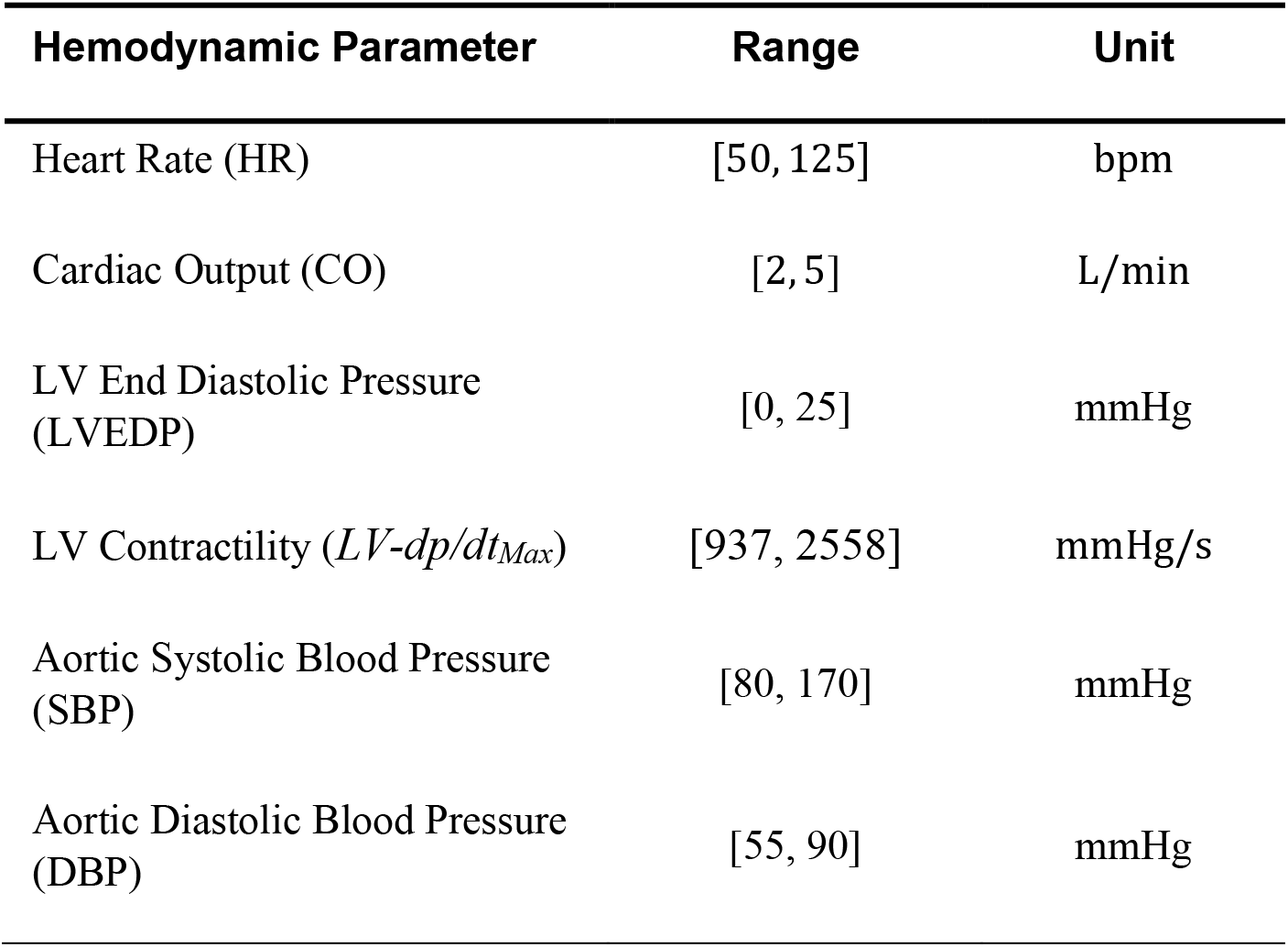
Summary of the hemodynamic conditions in the in-vitro experimentations.

### 2.4. Hemodynamic analysis

In this work, we employ both aortic input impedance as well as wave intensity (WI) as metrics to examine the dynamical and physiological accuracy of our proposed experimental setup.

#### Aortic input impedance

Aortic input impedance in pulsatile flow is defined as the ratio of pressure to flow in the frequency domain. It is a measure of the amount by which the aorta ‘impedes’ the flow. Input impedance is an index of arterial properties and is given by the expression [28, 32]

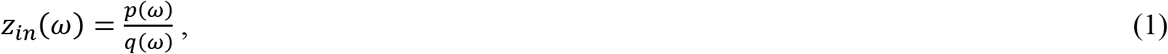

where *ω* is the frequency, *z_in_*(*ω*) is input impedance, *p*(*ω*) and *q*(*ω*) are the Fourier representations of the measured pressure and flow computed through applications of the fast Fourier transform (using *fft* in Matlab (Mathworks inc)).

#### Wave intensity

WI analysis is a well-established pulse wave analysis technique for quantifying the energy carried by arterial waves [33]. WI is determined by incremental changes in pressure and flow velocity, and hence requires measurements of both. A typical pattern of WI consists of a large amplitude forward (positive) peak (corresponding to the initial compression caused by a left ventricular contraction) followed by a small amplitude backward (negative) peak (corresponding to reflections from the initial contraction) which itself may be followed by a moderate amplitude forward decompression wave [33, 34]. The WI, called *dI*(*t*), is defined by the expression

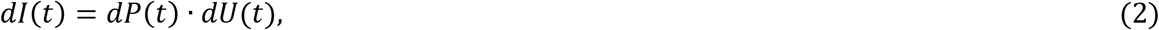

where *t* is the time, *P*(*t*) is the measured pressure, and *U*(*t*) is the measured velocity. For acquiring simultaneous measurements of pressure and flow for quantifying both impedance and WI, we use a Millar pressure system and Transonic flow meter as described in Section 2.3.

## 3. Results

### 3.1. Physiological accuracy of the atrioventricular-aortic hemodynamics

Figure 5 presents pressure waveforms at the LA, the LV, the ascending aorta, and the abdominal aorta corresponding to a baseline case of a healthy patient (i.e., *HR*=75 bpm and *CO*=5 L/min). A mid-range aorta (Aorta-3 of Table 2) is used for this measurement. The baseline LVEDP is 7 mmHg and the computed TPR is 19.32 mmHg.min/L, both of which are consistent with a normal healthy patient [28, 30–32].

**Figure 5.**
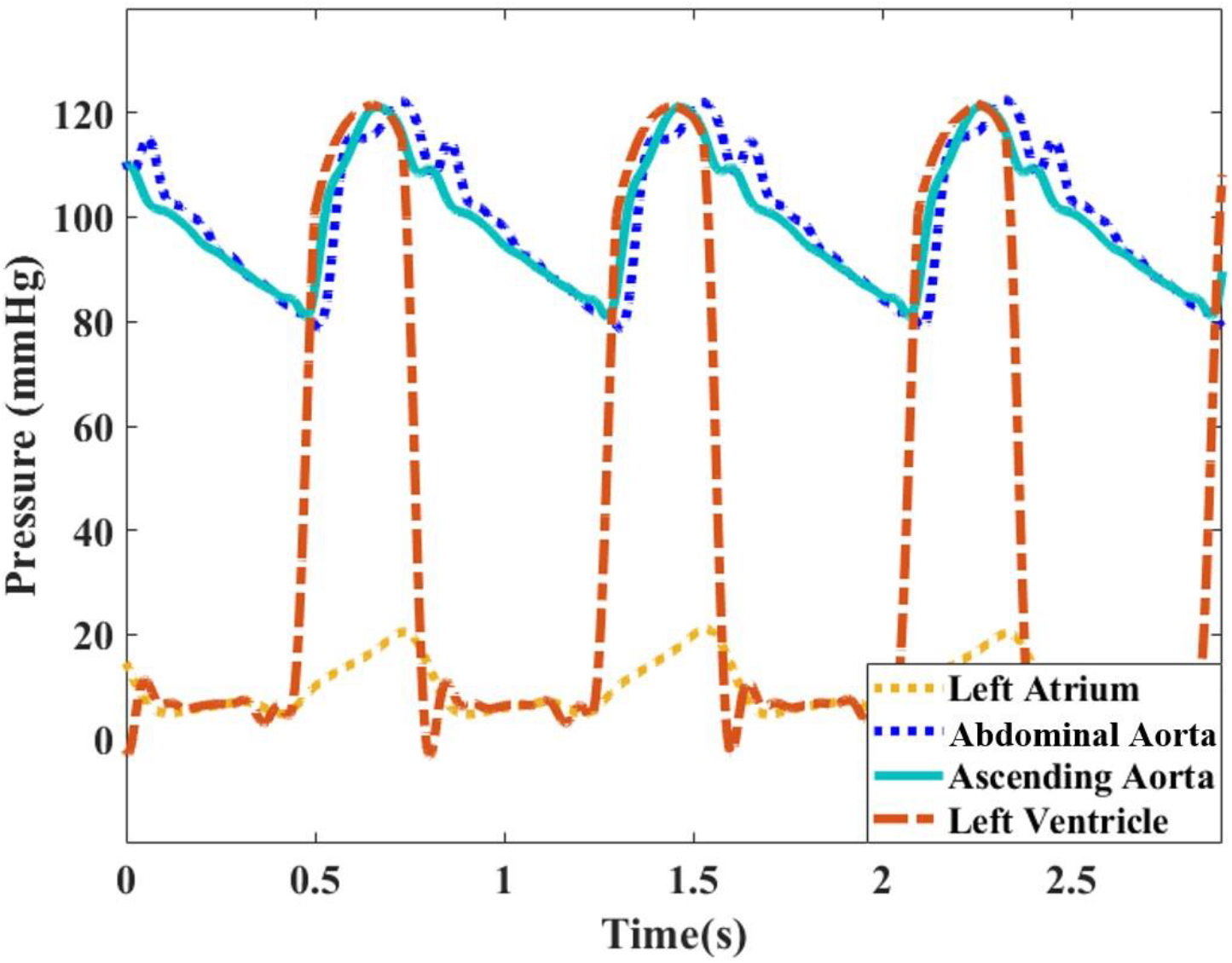
Pressure waveforms at different anatomical sites including the left atrium, the left ventricle, the ascending aorta, and the abdominal aorta. Waveforms are plotted for a normal healthy case (Aorta-3 of Table 2) corresponding to *CO* = 5 L/min and *HR* = 75 bpm

Figure 6 presents simultaneous pressure and flow measurements at the ascending aorta for different cardiac outputs (i.e., *CO* = 2, 3, 4, 5 L/min). For these measurements, Aorta-3 and HR=75 bpm are again employed. The expected fiducial features of the pressure and flow waveforms, including the pressure dicrotic notch, pressure augmentation, and the valve closure effect (zero flow during the diastole), can be observed in Figure 6. Results demonstrate that our setup is indeed sensitive to the increase in both aortic pressure and flow that arises from an increase in total cardiac output.

**Figure 6.**
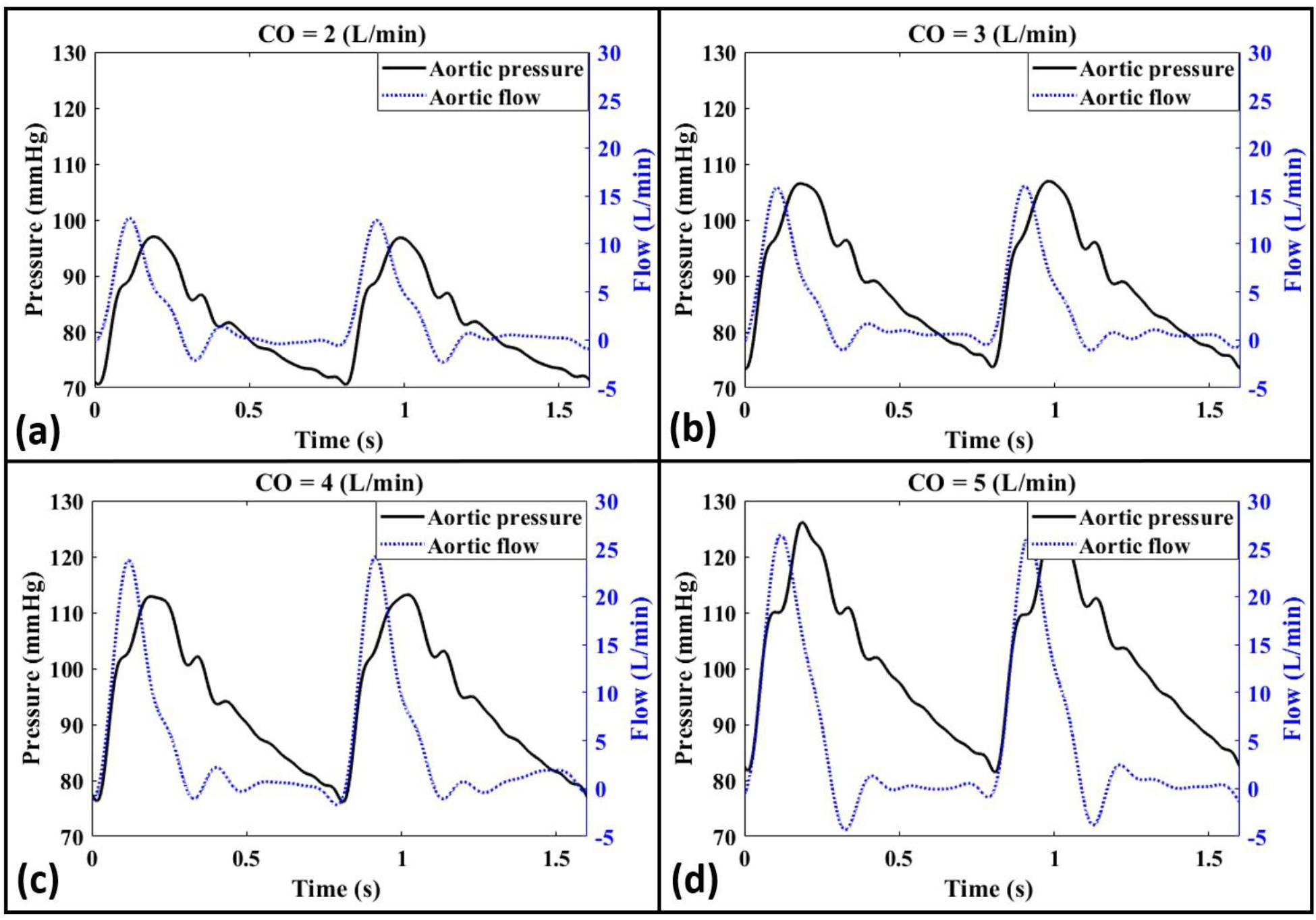
Demonstration of simultaneous aortic pressure and aortic flow measurements for Aorta-3 at different cardiac outputs corresponding to **(a)** *CO* = 2 L/min, **(b)** *CO* = 3 L/min, **(c)** *CO* = 4 L/min, and **(d)** *CO* = 5 L/min. Heart rate is 75 bpm for all the cases.

Figures 7a and 7b provide the corresponding WI (Eq. (2)) from the simultaneously-measured pressure and flow waveforms for different cardiac outputs and aortas, respectively. Typically-expected physiological patterns of WI are well-established in literature for clinical data [33]. Such patterns can be used as reference for assessing the physiological correctness of the WI profiles generated by our in-vitro setup and, indeed, our results appear consistent. Figures 7c and 7d additionally present the aortic input impedance for different cardiac outputs of a fixed aorta (Aorta-3) and for different aortas at the same cardiac output (CO=5 L/min), respectively. All measured data correspond to HR=75 bpm. The raw data corresponding to all the figures of this section is available online.

**Figure 7.**
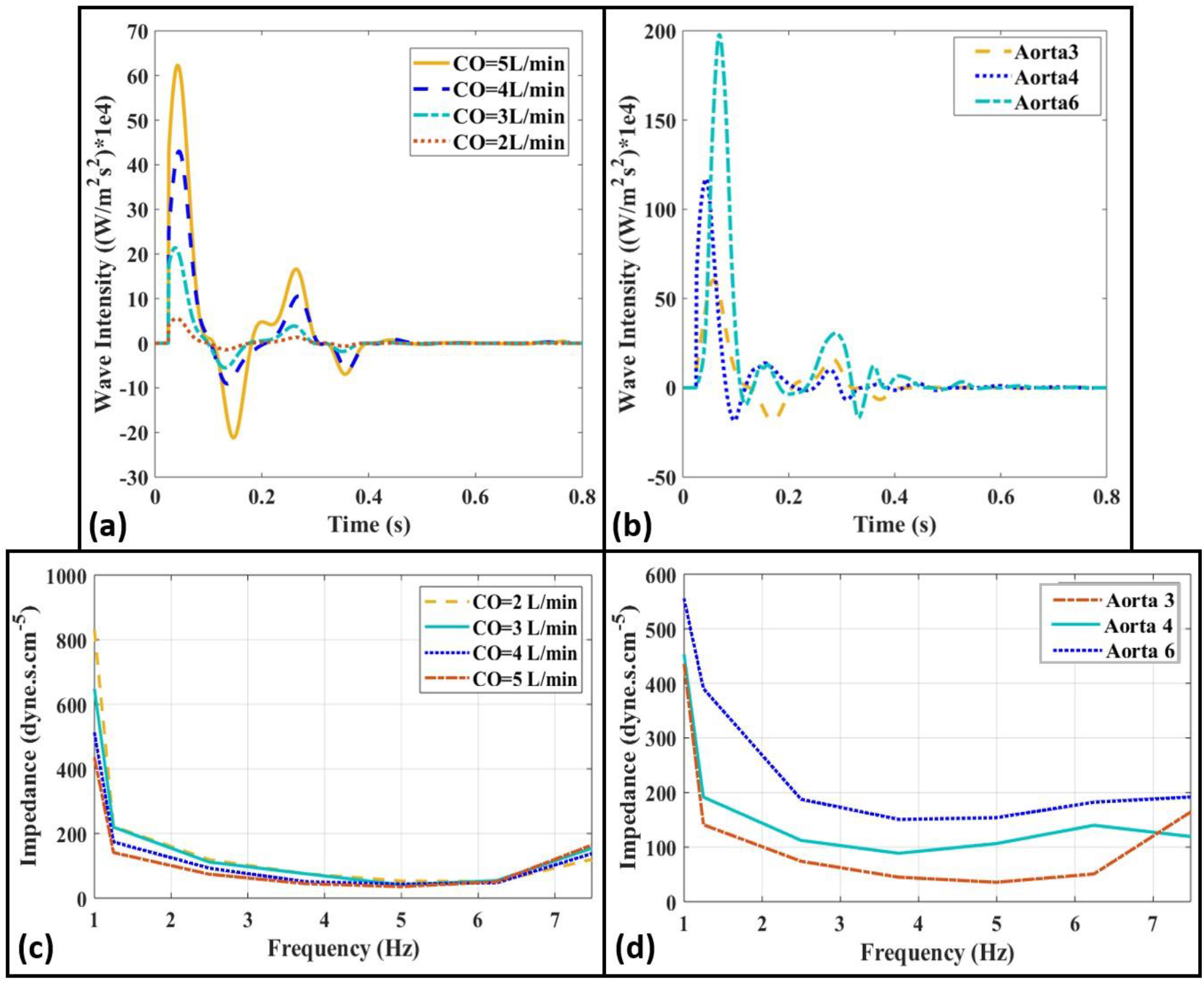
Wave intensity profiles measured at the ascending aorta for **(a)** different cardiac outputs of the same Aorta-3 (*CO* = 2,3,4,5 L/min), **(b)** different aortas for the same *CO* = 5 L/min (Aorta-3, 4, 6). **(c, d)** Aortic input impedances corresponding to the same configurations of **(a,b**). Heart rate is 75 bpm for all the cases.

### 3.2. Effects of cardiac function on coupled atrioventricular-aortic hemodynamics

In this section, we study the effects of cardiac function determinants (i.e., stroke volume, contractility, and heart rate) on measured pressure waveforms. Figure 8 presents such measured pressures at different anatomical sites for different stroke volumes corresponding to normal (*CO* = 5 L/min) and reduced cardiac outputs for simulating impaired cardiac function (i.e., *CO* = 2,3,4 L/min). Measurements are conducted for Aorta-3 at a fixed (normal) contractility and HR=75 bpm.

**Figure 8.**
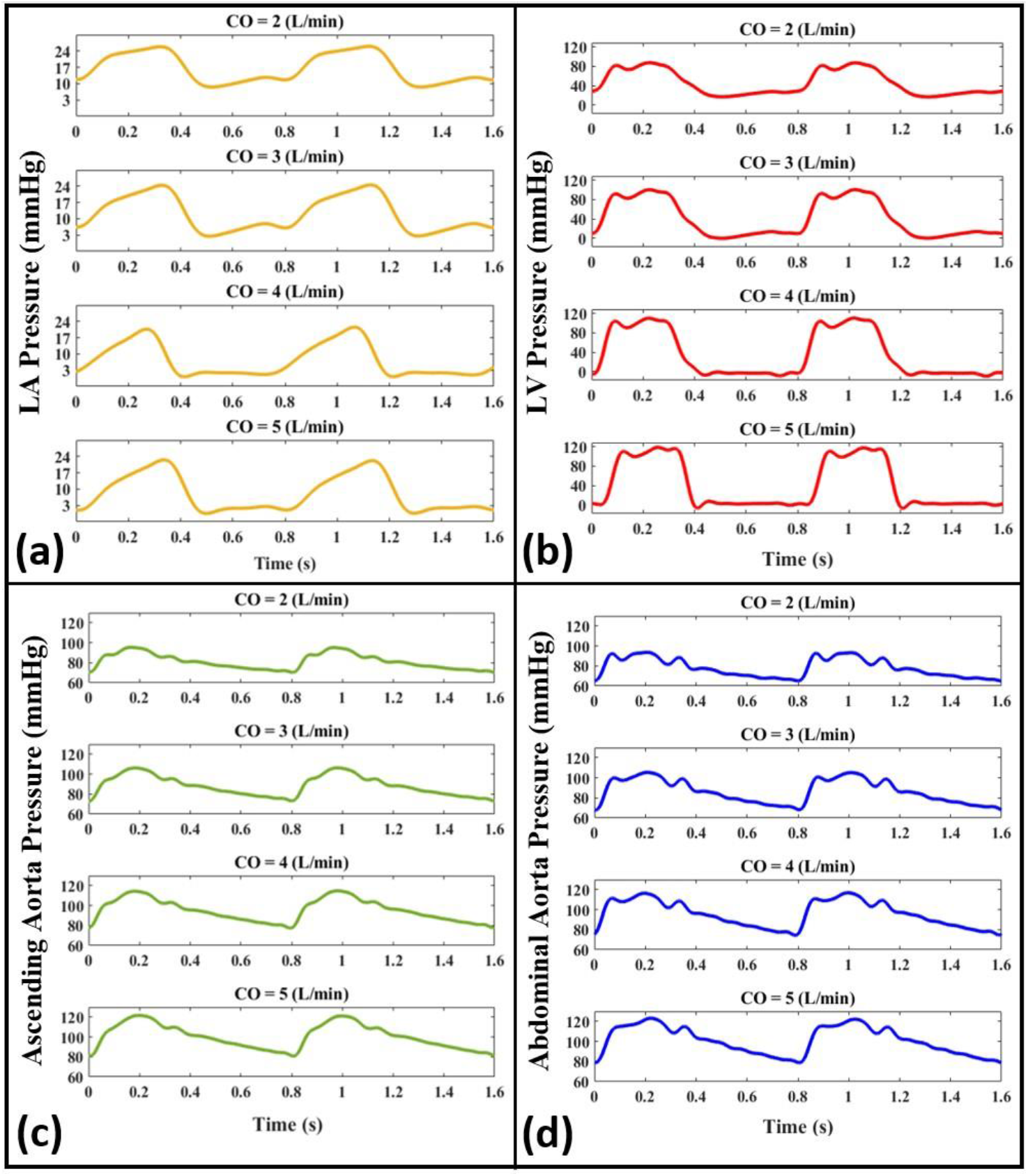
Pressure waveforms for normal (*CO* = 5 L/min) and impaired (*CO* = 2,3,4 L/min) conditions of cardiac function at different anatomical sites including **(a)** the left atrium, **(b)** the left ventricle, **(c)** the ascending aorta, and **(d)** the abdominal aorta. Waveforms are plotted for Aorta-3 at *HR* = 75 bpm and a fixed (normal) contractility.

Figure 9 presents measured pressure data at the LA, the LV, the ascending aorta, and the abdominal aorta for different contractility conditions, i.e., high contractility (LV-*dp/dt_Max_*=2558.3 mmHg/s), normal contractility (*LV-dp/dt_Max_*= 1432.7 mmHg/s), and low contractility (*LV-dp/dt_Max_*=937.4 mmHg/s) at a fixed HR of 75 bpm and CO of 5 L/min.

**Figure 9.**
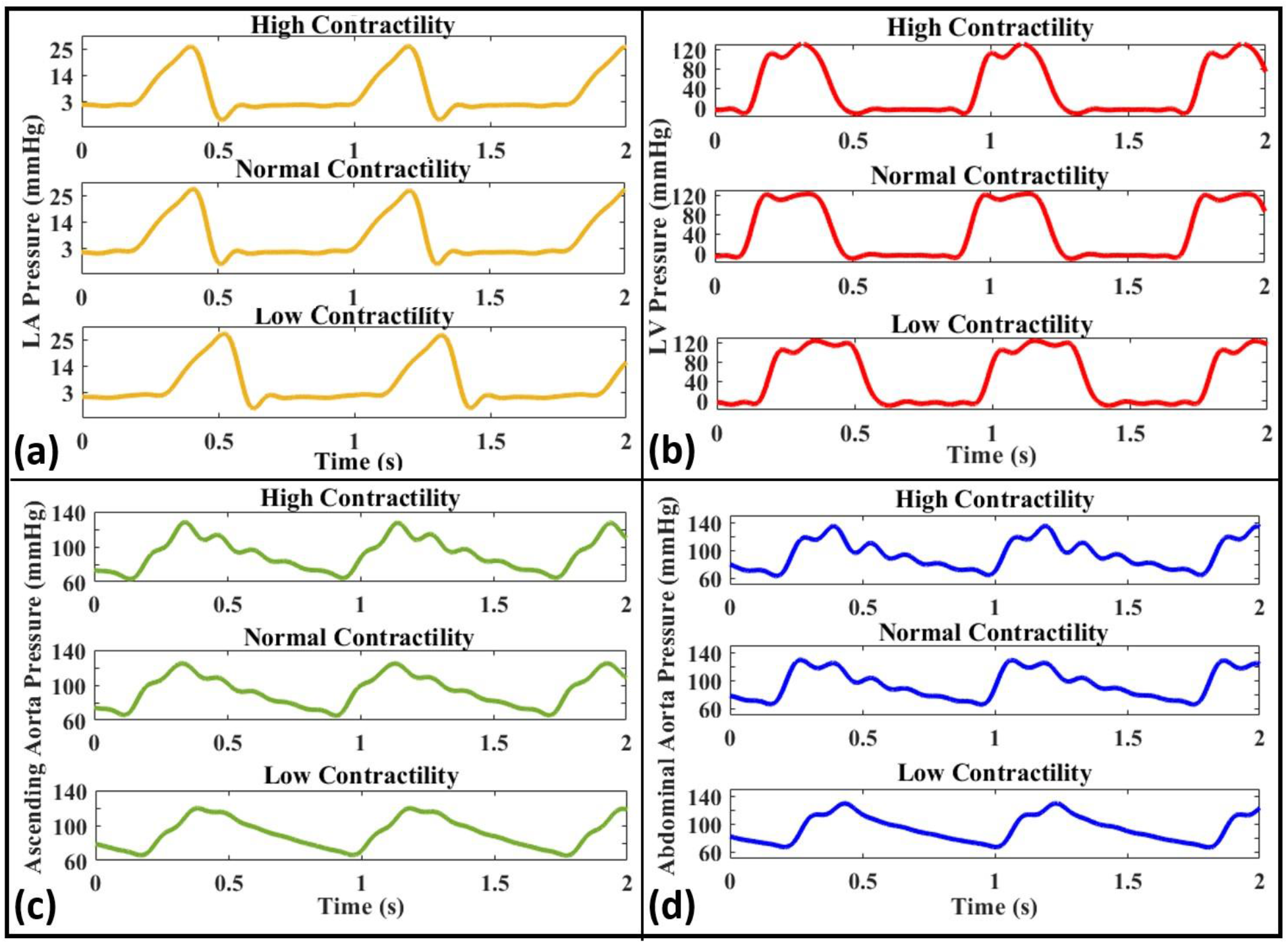
Pressure waveforms for different contractility conditions including high (LV-*dp/dt_Max_*=2558.3 mmHg/s), normal (*LV-dp/dt_Max_*= 1432.7 mmHg/s), and low contractility (LV-*dp/dt_Max_*=937.4 mmHg/s) at **(a)** the left atrium, **(b)** the left ventricle, **(c)** the ascending aorta, and **(d)** the abdominal aorta (Aorta-4, HR = 75 bpm, CO = 5 L/min).

Examples of measured pressure waveforms for different heart rates (*HR* = 50, 75, 100, 125 bpm) at the LA and the ascending aorta are presented in Fig. 10 for a fixed (normal) contractility condition. These waveforms are measured using Aorta-3 at *CO* = 5 L/min. The raw data corresponding to all figures of this section is available online.

**Figure 10.**
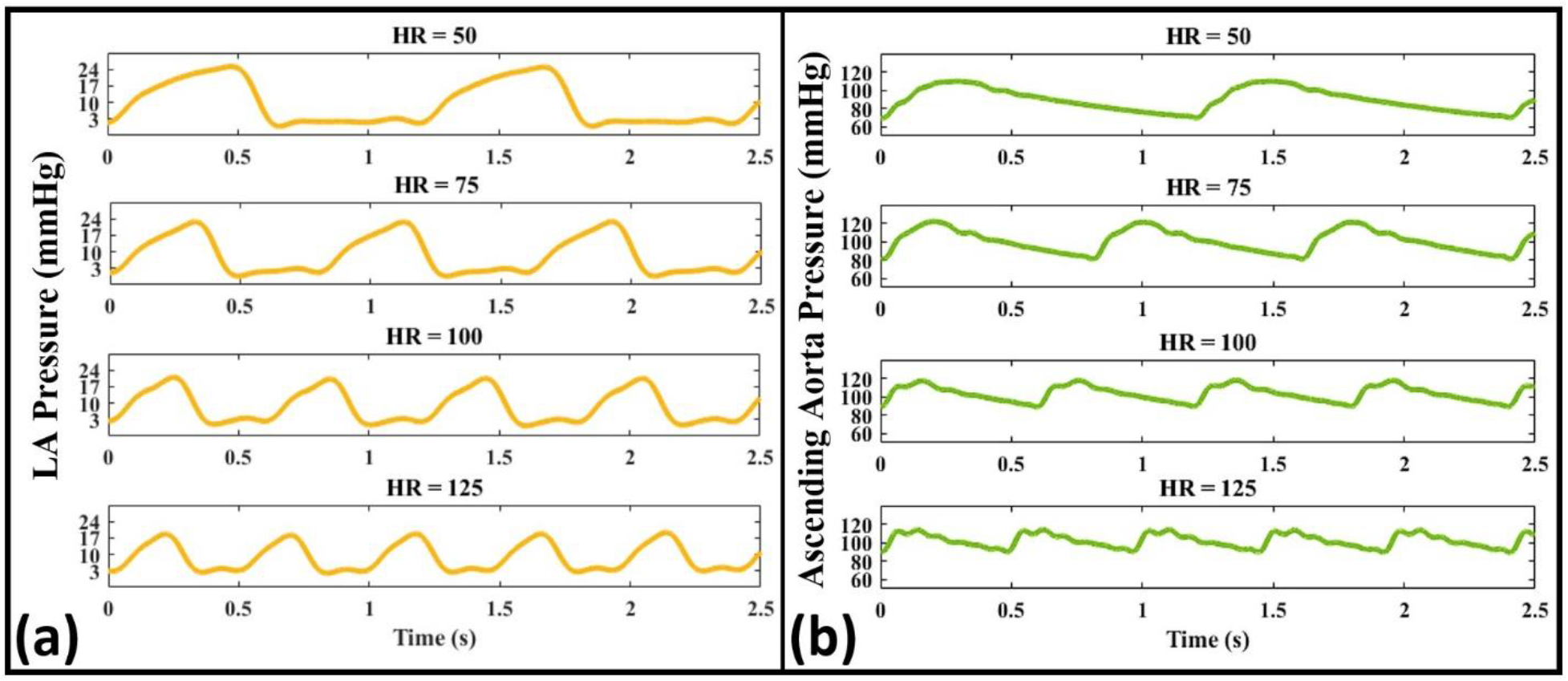
Pressure waveforms for different heart rates (*HR* = 50, 75,100,125 bpm) at **(a)** the left atrium and **(b)** the ascending aorta (Aorta-3, *CO* = 5 L/min).

### 3.3. Effects of afterload on coupled atrioventricular-aortic hemodynamics

In this section, we study using our setup the effects on hemodynamic waves of peripheral resistance and aortic compliance (determinants of the afterload). Figure 11 demonstrates the effect of high peripheral resistance on the measured pressure waveforms inside the LA, the LV, the ascending aorta, and the abdominal aorta. This effect is modeled using the terminal clamps mounted at the outlets of the artificial aorta (as described in Section 2). All data is from Aorta-3 and is simulated at HR=75 bpm.

**Figure 11.**
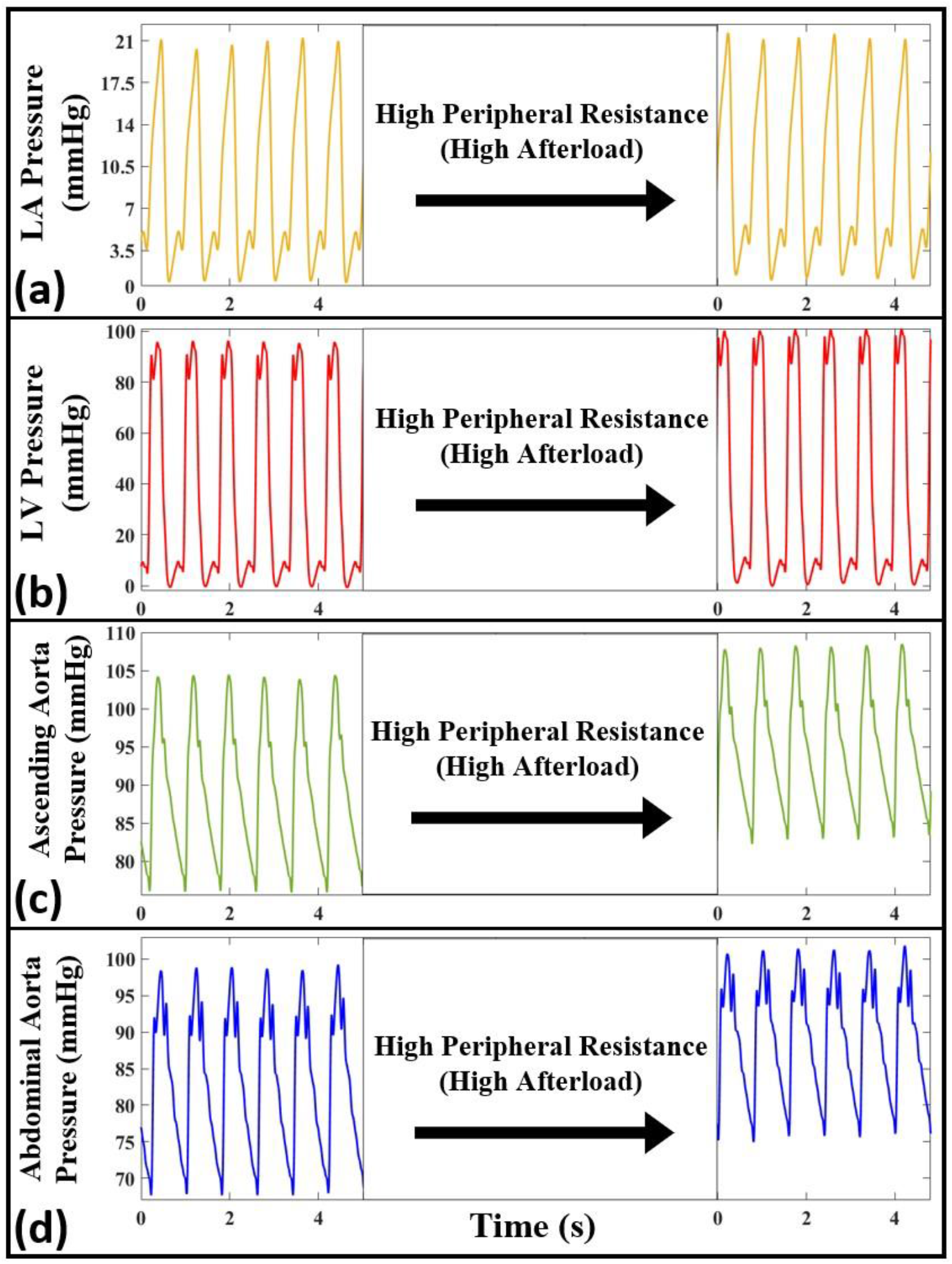
The measured pressure waveforms for normal and high peripheral resistance (i.e., high afterload) conditions at different anatomical sites including (a) the left atrium, (b) the left ventricle, (c) the ascending aorta, and (d) the abdominal aorta. Waveforms are produced from Aorta-3 and correspond to *HR* = 75 bpm.

Figure 12 presents pressure waveforms measured at different anatomical sites for eight different aortas representing various aortic compliances (see Table 2 for their characteristics). Measured data corresponds to HR=75 bpm and CO=5 L/min. The raw data corresponding to all figures of this section is available online.

**Figure 12.**
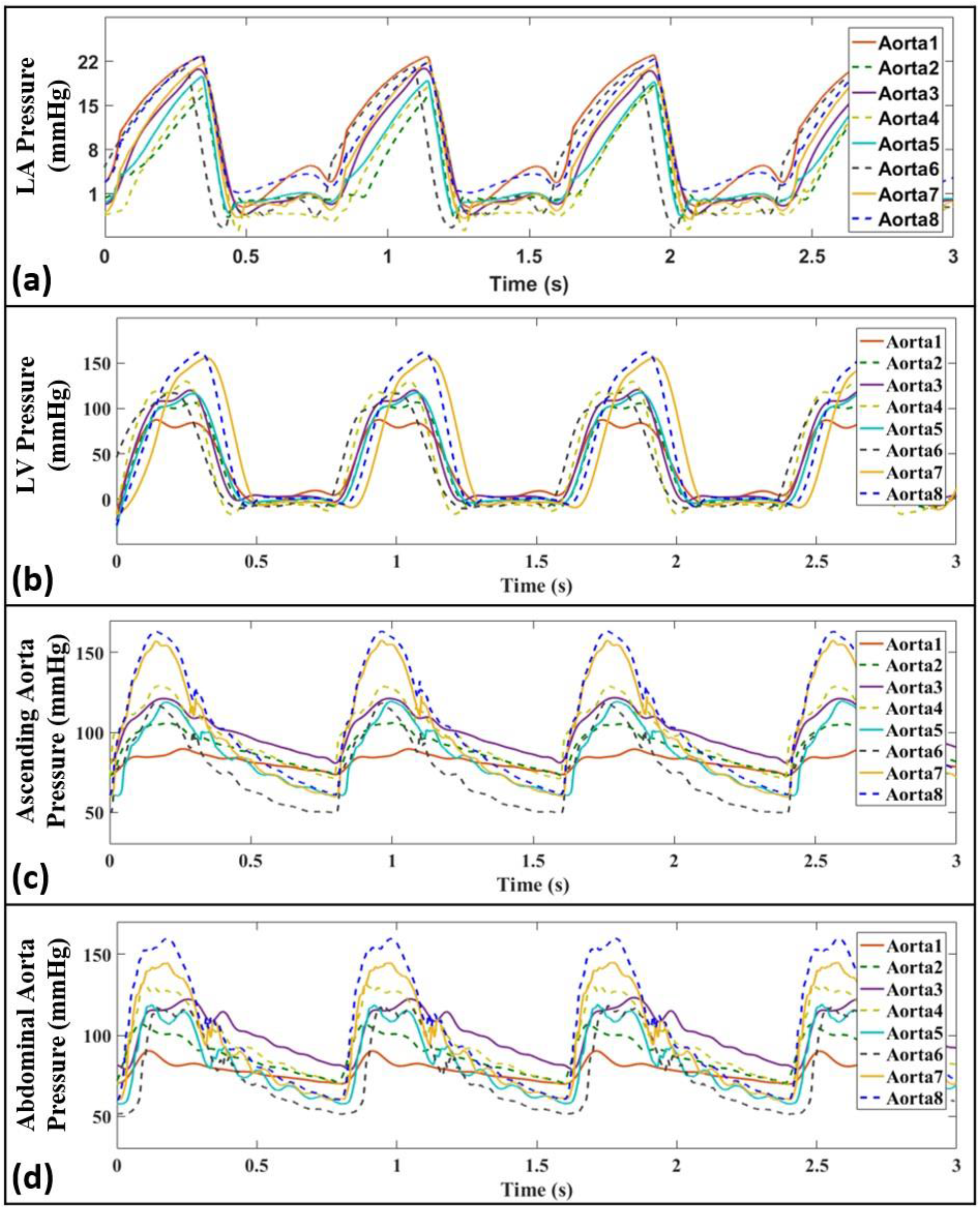
Pressure waveforms, corresponding to different aortas (i.e., Aorta-1 to 8) representing various aortic compliances, at different anatomical sites including **(a)** the left atrium, **(b)** the left ventricle, **(c)** the ascending aorta, and **(d)** the abdominal aorta. Waveforms correspond to *CO* = 5 L/min and *HR* = 75 bpm.

### 3.4. Effects of cardiac preload on coupled atrioventricular-aortic hemodynamics

Figure 13 demonstrates the effects of reduced and increased preload (relative to the baseline condition of Fig. 5) on pressure waveforms measured at four different anatomical sites. The reduced and increased preload cases correspond to values of LVEDP corresponding to 0 mmHg and of 25 mmH*g*, respectively. The same heart rate (HR = 75 bpm), artificial aorta (Aorta-3), and contractility condition (normal) are considered for both measurements. The raw data corresponding to Fig. 13 is available online.

**Figure 13.**
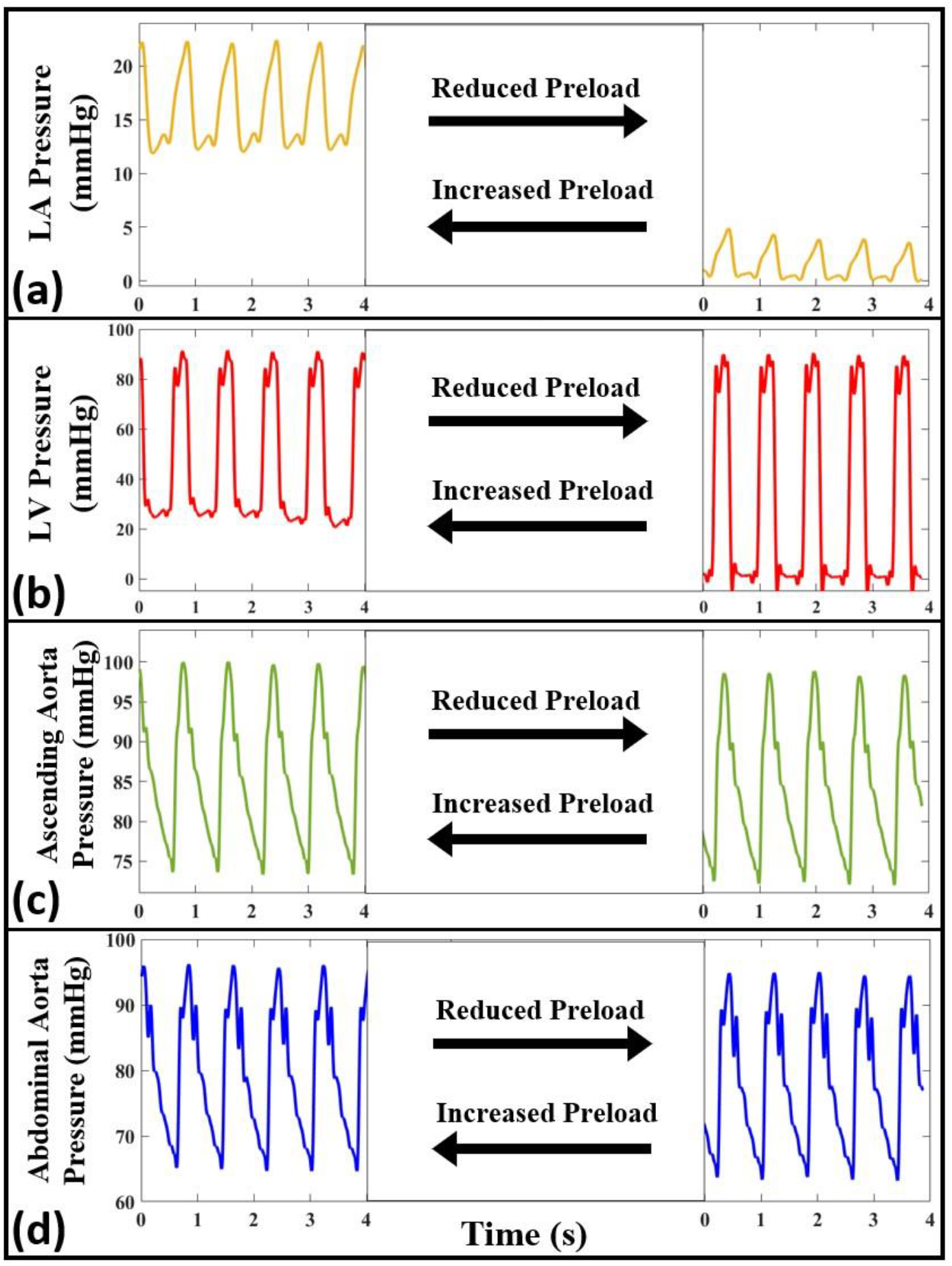
The measured pressure waveforms for increased and reduced preload conditions at different anatomical sites including **(a)** the left atrium, **(b)** the left ventricle, **(c)** the ascending aorta, and **(d)** the abdominal aorta. Waveforms are measured in Aorta-3 and correspond to HR = 75 bpm.

### 3.5. Applicability of the setup for non-invasive measurements

Figures 14a and 14b present pressure waveform measurements using a high-fidelity Piezo-tip Millar pressure catheter (an invasive measurement) and an FDA-approved non-invasive handheld device (called Vivio [29]), respectively. Figure 14b shows a sample screenshot of the Vivio iPad application, illustrating the heart sound measured using Vivio. These measurements are conducted on Aorta-4 at CO=4.5 L/min and HR=75 bpm.

**Figure 14.**
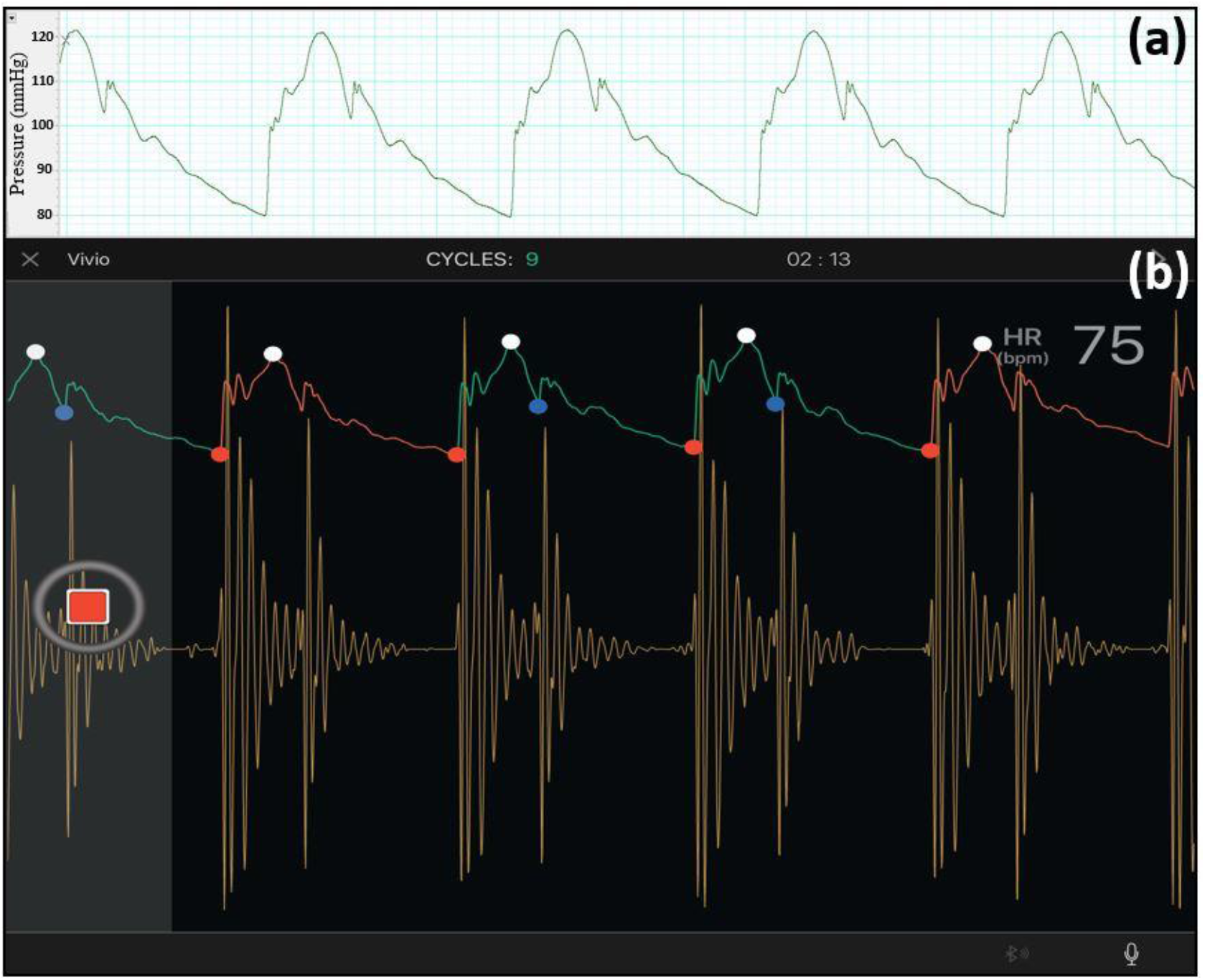
Example of simultaneous invasive and noninvasive measurements using **(a)** a Piezo-tip Millar MIKRO-TIP catheter transducer (each square accounts for 50 milliseconds), and **(b)** Vivio (a wireless optical tonometer [29]). Data is recorded in Aorta-4 at CO=4.5 L/min and HR=75 bpm. The Vivio system simultaneously measures pulse pressure and heart sound, communicating with a tablet or an iPad for real-time data display and for capturing via BLE.

## 4. Discussion

In this paper, we have introduced and described a state-of-the-art experimental model for the in-vitro investigation of hemodynamics and wave propagation in the systemic circulation. Results demonstrate the ability of our in-vitro experimental setup to reproduce physiological functions of the LV, the LA, and the vascular system that correspond to both healthy and diseased conditions. These functions have been evaluated across a wide range of hemodynamic conditions (i.e., different afterload, preload, and contractility).

For assessing the physiological relevancy of the setup, we have measured the pressure waveforms at different anatomical sites of the system including inside the LV, inside the LA, the ascending aorta and the abdominal aorta (Fig. 5). For the baseline pressure measurements, the LV- excitation pump has been operated at 75 bpm with a measured CO of 5 L/min (corresponding to normal contractility). As can be observed in Fig. 5 and Fig. 6, our system is able to generate the main physiological features of the pressure and flow waveforms as found in the human cardiovascular system. These include: *i*) the physiological development of the pressure inside the LV and the aorta during systole (corresponding to the LV-arterial coupling), *ii*) the presence of the dicrotic notch due to the aortic valve closure (corresponding to the LV-arterial decoupling), *iii*) the interaction of the ventricle and the atrium during the filling phase of the LV via the mechanical mitral valve (corresponding to the LV-LA coupling and decoupling), *iv*) the increase in the pulse pressure as the wave propagates downstream towards the abdominal aorta, and *v*) the physiological point-to-point consistency of the pressure and flow [28, 30–32]. Indeed, Figs. 5 and 6 demonstrate that our setup generates physiologically-relevant waveform morphologies [28, 30–32]. The computed augmentation index (*AI*, the percentage of the augmented pressure in the central pulse pressure) and form factor (*FF*, the ratio between mean pulse pressure and pulse pressure) [28] are within physiological ranges (*AI*=29.8% and *FF*=0.33). For the baseline measurements, the mean aortic pressure corresponds to 94.25 mmHg, the aortic SBP to 121.20 mmHg, the aortic DBP to 80.78 mmHg, and the total peripheral resistance (TPR) to 18.85 mmHg·min/L. In order to evaluate the dynamic accuracy of the setup, we have also considered WI (Figs. 7a and 7b), which is an energy-based clinical index quantifying the power carried in the arterial system (Section 2.4). The three major peaks of WI are successfully generated by our in-vitro setup for different cardiac outputs and aortas. We have shown in Fig. 7a that as CO decreases, so does the amplitude of all three major peaks as expected. We have also employed another measure, aortic input impedance [32, 35], for similarly assessing the dynamic accuracy of our system. As demonstrated in Figs. 7c and 7d, our measured impedances fall well-within the expected physiological ranges and profiles [17, 32].

We have further studied the effects of cardiac function on the measurements of our coupled in-vitro system. At a fixed cardiac output and LV contractility, Fig. 10 demonstrates that higher heart rates lead to decreased stroke volumes and consequently result in lower pulse pressures inside the aorta and the atrium (a result expected from previous studies [4, 10]). Additionally, at a fixed CO and HR, we have investigated the impact of LV contractility alone, achieved by changes in the input profile of the LV-excitation pump, on the propagated pressure waveforms. Our measured data (Fig. 9) indicates that impaired contractility, independently of the stroke volume, results in lower pulse pressure in the ascending and abdominal aorta as well as results in changes to waveform morphology. Such a result is in agreement with recent clinical literature [36, 37]. We have also further quantified the effects of changing LV contractility on the ascending aorta pressure waveform by calculating the value of *dp/dt_Max_* for such waveforms. The *dp/dt_Max_* values of the ascending aorta have been determined as *dp/dt_Max_*=443.4 mmHg/s for high contractility, *dp/dt_Max_*=404.8 mmHg/s for normal contractility, and *dp/dt_Max_*=358.0 mmHg/s for low contractility. The calculated aortic *dp/dt_Max_* values verify the proper LV-arterial coupling at which different LV contractility conditions physiologically affect the propagated wave toward the arterial system in our in-vitro setup.

Our in-vitro setup is also a valuable tool for simulating cardiovascular diseases related to impaired cardiac function. We have successfully simulated physiological systolic dysfunction (from mild to severe) for different levels of cardiac output (Fig. 8). The hemodynamic results from the experimental setup demonstrate that reduced cardiac output leads to lower mean and pulse pressures inside the LV, the ascending aorta and the abdominal aorta. However, reduced cardiac output results in higher pressures inside the LA as well as higher LVEDP values. This is in agreement with clinical observations suggesting that the occurrence of heart failure is highly correlated with elevated LVEDP [30, 38].

Previous studies have demonstrated that aortic compliance is a contributor of the pulsatile load on the global cardiovascular system and that the peripheral resistance is a contributor to the steady portion [25]. In this manuscript, we have presented studies on both these afterloads using our in-vitro setup, and their effects on in-vitro hemodynamics have been investigated. As supported by Fig. 12, our in-vitro setup is able to capture the effects of aortic compliance on the ventricular and arterial pulse pressures. As expected [4, 10, 31], lower aortic compliances (larger stiffnesses) lead to higher ventricular and arterial pressures. Figure 11 presents our in-vitro simulations for increased peripheral resistance where it can be seen that, in addition to the expected rise in the mean pressure in the ascending and abdominal aorta, our setup is also able to capture the increase in ventricular pressure that arises from increased peripheral resistance (a demonstration that our design faithfully considers direct LV-aortic coupling). Another effect of increasing the peripheral resistance in our setup is the drop in CO from 4 L/min to 3.3 L/min. This demonstrates successful direct hemodynamic coupling of the LV sac and the aorta. It also demonstrates that the general pumping behavior in our system acts exactly similarly to the pumping characteristics of the heart (which is neither flow nor pressure source [3, 4]).

The effects of cardiac preload (a major contributor to the cardiovascular system) on the coupled atrioventricular-aortic hemodynamics has been presented in Fig. 13. As expected, the LA pressure and the diastolic part of the LV pressure increase significantly after increasing the cardiac preload (LVEDP increases by 25 mmHg) [30, 38]. This effect illustrates the coupling characteristics between the LA and LV in our setup. Such results point to the physiological relevancy of our in-vitro coupled atrioventricular-aortic simulator for studying LV-diastolic dysfunction.

Lastly, we have examined the applicability of our setup for non-invasive hemodynamic measurements. Non-invasive measurement methodologies are of particular interest due to currently-in-development wearable technologies for monitoring and diagnostic purposes. We have compared simultaneous invasive and non-invasive measurements using our setup, and have demonstrated (Fig. 14) that the non-invasive waveforms measured by Vivio [29] successfully represent morphological and physiological features (e.g., the pressure augmentation point or the dicrotic notch) of the corresponding invasive waveforms Additionally, as can be observed in Fig. 14b, both the first as well as the second heart sounds are successfully generated by our constructed setup. Our non-invasive measurements are also physiological and demonstrate consistent phonocardiogram waveforms [39, 40].

### 4.1. Limitations and future work

Previous studies have shown that the non-Newtonian behavior of blood impacts the hemodynamics of the cardiovascular system [41, 42]. However, it is well-established that the fluid viscosity has negligible effects on hemodynamic waves [4, 32].

Future work includes developing a large in-vitro database using our setup for resolving issues related to usually small-sized in-vivo databases in order to facilitate specific hemodynamics-based big-data studies [43–45]. In addition, the setup can be used to simulate more complicated diseases, e.g., myocardial infarction, hemorrhagic shock, or aortic coarctation, where several physiological responses need to be simulated (i.e., changing our setup based on the main pathophysiology/abnormalities of the corresponding disease condition) [30, 46, 47].

## 5. Conclusions

The design and fabrication of an in-vitro cardiovascular simulator are presented in this study. We have demonstrated the physiological relevancy of the setup by studying the expected physiological features in the measured hemodynamic waveforms. Our setup can be used for modeling the effects of major contributors of the cardiovascular system such as LV contractility, afterload, and cardiac preload. Studying such effects ultimately improves understanding of the underlying hemodynamic mechanisms of different cardiovascular diseases. Additionally, due to no media change between our LV and aorta, non-invasive sound and pulse measurements are also available via the setup; hence our system can be employed for assessing the accuracy of any newly-developed non-invasive methodology. Our proposed setup is also useful for cardiovascular big-data studies, potentially providing large in-vitro databases for use in deep learning or machine learning studies. Ultimately, this setup is a practical in-vitro tool for preliminary cardiovascular studies prior to expensive and high-risk in-vivo trials.

## Supporting information

**S1 File.** Example of a physiological input waveform for the piston pump displacement.

**S2 File.** The detailed CAD design of the constructed setup (3D CAD file).

## Acknowledgments

The authors would like to thank Dr. Faisal Amlani for productive discussion.

## Author contributions

*RA*: Data collection, manuscript preparation, concept/design, setup fabrication, vessel fabrication, data analysis/interpretation, results, and figures

*AA*: Data collection, manuscript preparation, vessel fabrication, data analysis/interpretation, results, and figures

*HW*: Data collection, manuscript preparation, vessel fabrication, data analysis/interpretation, results, and figures

*SN*: Data collection, manuscript preparation, vessel fabrication, data analysis/interpretation, results, and figures

*SW*: Concept/design, setup fabrication

*NMP*: Research supervision, concept/design, manuscript proofread, data analysis/interpretation

## References

1. Starling MR. Left ventricular-arterial coupling relations in the normal human heart. American heart journal. 1993;125(6):1659–66.

2. Bell V, Mitchell GF. Influence of vascular function and pulsatile hemodynamics on cardiac function. Current hypertension reports. 2015;17(9):1–9.

3. Pahlevan NM, Gharib M. Aortic wave dynamics and its influence on left ventricular workload. PloS one. 2011;6(8):e23106.

4. Pahlevan NM, Gharib M. A bio-inspired approach for the reduction of left ventricular workload. PLoS One. 2014;9(1):e87122.

5. Mitchell GF. Effects of central arterial aging on the structure and function of the peripheral vasculature: implications for end-organ damage. Journal of applied physiology. 2008;105(5):1652–60.

6. Mitchell GF, Tardif J-C, Arnold JMO, Marchiori G, O’Brien TX, Dunlap ME, et al. Pulsatile hemodynamics in congestive heart failure. Hypertension. 2001;38(6):1433–9.

7. Bell V, McCabe EL, Larson MG, Rong J, Merz AA, Osypiuk E, et al. Relations between aortic stiffness and left ventricular mechanical function in the community. Journal of the American Heart Association. 2017;6(1):e004903.

8. Mitchell GF, van Buchem MA, Sigurdsson S, Gotal JD, Jonsdottir MK, Kjartansson Ó, et al. Arterial stiffness, pressure and flow pulsatility and brain structure and function: the Age, Gene/Environment Susceptibility–Reykjavik study. Brain. 2011;134(11):3398–407.

9. Ooi H, Chung W, Biolo A. Arterial stiffness and vascular load in heart failure. Congestive heart failure. 2008;14(1):31–6.

10. O’Rourke MF. Steady and pulsatile energy losses in the systemic circulation under normal conditions and in simulated arterial disease. Cardiovascular research. 1967;1(4):313–26.

11. Lam CS, Gona P, Larson MG, Aragam J, Lee DS, Mitchell GF, et al. Aortic root remodeling and risk of heart failure in the Framingham Heart study. JACC: Heart Failure. 2013;1(1):79–83.

12. Laskey WK, Kussmaul WG. Arterial wave reflection in heart failure. Circulation. 1987;75(4):711–22.

13. Pahlevan NM, Gharib M. In-vitro investigation of a potential wave pumping effect in human aorta. Journal of biomechanics. 2013;46(13):2122–9.

14. Cooper LL, Rong J, Benjamin EJ, Larson MG, Levy D, Vita JA, et al. Components of hemodynamic load and cardiovascular events: the Framingham Heart Study. Circulation. 2015;131(4):354–61.

15. Cooper LL, Rong J, Pahlevan NM, Rinderknecht DG, Benjamin EJ, Hamburg NM, et al. Intrinsic frequencies of carotid pressure waveforms predict heart failure events: the framingham heart study. Hypertension. 2021;77(2):338–46.

16. Gaddum N, Alastruey J, Chowienczyk P, Rutten MC, Segers P, Schaeffter T. Relative contributions from the ventricle and arterial tree to arterial pressure and its amplification: an experimental study. American Journal of Physiology-Heart and Circulatory Physiology. 2017;313(3):H558–H67.

17. Segers P, Dubois F, De Wachter D, Verdonck P. Role and relevancy of a cardiovascular simulator. Cardiovascular Engineering. 1998;3:48–56.

18. Timms D, Hayne M, McNeil K, Galbraith A. A complete mock circulation loop for the evaluation of left, right, and biventricular assist devices. Artificial organs. 2005;29(7):564–72.

19. Cox C, Najjari MR, Plesniak MW. Three-dimensional vortical structures and wall shear stress in a curved artery model. Physics of Fluids. 2019;31(12):121903.

20. Brindise MC, Chiastra C, Burzotta F, Migliavacca F, Vlachos PP. Hemodynamics of stent implantation procedures in coronary bifurcations: an in vitro study. Annals of biomedical engineering. 2017;45(3):542–53.

21. Peterson SD, Plesniak MW. The influence of inlet velocity profile and secondary flow on pulsatile flow in a model artery with stenosis. Journal of fluid mechanics. 2008;616:263–301.

22. Alastruey J, Khir AW, Matthys KS, Segers P, Sherwin SJ, Verdonck PR, et al. Pulse wave propagation in a model human arterial network: assessment of 1-D visco-elastic simulations against in vitro measurements. Journal of biomechanics. 2011;44(12):2250–8.

23. Alastruey J, Parker K, Peiró J, Byrd S, Sherwin S. Modelling the circle of Willis to assess the effects of anatomical variations and occlusions on cerebral flows. Journal of biomechanics. 2007;40(8):1794–805.

24. Aghilinejad A, Amlani F, King KS, Pahlevan NM. Dynamic effects of aortic arch stiffening on pulsatile energy transmission to cerebral vasculature as a determinant of brain-heart coupling. Scientific reports. 2020;10(1):1–12.

25. Mitchell GF. Aortic stiffness, pressure and flow pulsatility, and target organ damage. Journal of Applied Physiology. 2018;125(12):1871–80.

26. Cappon F, Wu T, Papaioannou T, Du X, Hsu P-L, Khir AW. Mock circulatory loops used for testing cardiac assist devices: A review of computational and experimental models. The International Journal of Artificial Organs. 2021;44(11):793–806.

27. Knoops PG, Biglino G, Hughes AD, Parker KH, Xu L, Schievano S, et al. A mock circulatory system incorporating a compliant 3D-printed anatomical model to investigate pulmonary hemodynamics. Artificial organs. 2017;41(7):637–46.

28. Salvi P. Pulse waves. How vascular hemodynamics affects Blood pressure. 2012.

29. Rinderknecht D, De Balasy JM, Pahlevan NM. A wireless optical handheld device for carotid waveform measurement and its validation in a clinical study. Physiological measurement. 2020;41(5):055008.

30. Ahrens T, Taylor LA. Hemodynamic waveform analysis: WB Saunders Company; 1992.

31. Fung Y-c. Biomechanics: circulation: Springer Science & Business Media; 2013.

32. Zamir M. Hemo-dynamics: Springer; 2016.

33. Aghilinejad A, Amlani F, Liu J, Pahlevan NM. Accuracy and applicability of non-invasive evaluation of aortic wave intensity using only pressure waveforms in humans. Physiological Measurement. 2021;42(10):105003.

34. Kang J, Aghilinejad A, Pahlevan NM. On the accuracy of displacement-based wave intensity analysis: Effect of vessel wall viscoelasticity and nonlinearity. PloS one. 2019;14(11):e0224390.

35. Zamir M. Mechanics of blood supply to the heart: wave reflection effects in a right coronary artery. Proceedings of the Royal Society of London Series B: Biological Sciences. 1998;265(1394):439–44.

36. Fok H, Guilcher A, Li Y, Brett S, Shah A, Clapp B, et al. Augmentation pressure is influenced by ventricular contractility/relaxation dynamics: novel mechanism of reduction of pulse pressure by nitrates. Hypertension. 2014;63(5):1050–5.

37. Kim HK, Pinsky MR. Effect of tidal volume, sampling duration, and cardiac contractility on pulse pressure and stroke volume variation during positive-pressure ventilation. Critical care medicine. 2008;36(10):2858.

38. Iskandrian A, Segal B, Hamid HAKKI A. Left ventricular end-diastolic pressure in evaluating left ventricular function. Clinical cardiology. 1981;4(1):28–33.

39. Singh M, Cheema A. Heart sounds classification using feature extraction of phonocardiography signal. International Journal of Computer Applications. 2013;77(4).

40. Kelly R. Non-invasive registration of the arterial pressure pulse waveform using high-fidelity applanation tonometry. J Vasc Med Biol. 1989;1:142–9.

41. Bilgi C, Atalık K. Effects of blood viscoelasticity on pulsatile hemodynamics in arterial aneurysms. Journal of Non-Newtonian Fluid Mechanics. 2020;279:104263.

42. Wei H, Cheng AL, Pahlevan NM. On the significance of blood flow shear-rate-dependency in modeling of Fontan hemodynamics. European Journal of Mechanics-B/Fluids. 2020;84:1–14.

43. Alavi R, Dai W, Kloner RA, Pahlevan NM. A Hybrid Artificial Intelligence-Intrinsic Frequency Method for Instantaneous Determination of Myocardial Infarct Size. Circulation. 2020;142(Suppl_3):A15899–A.

44. Alavi R, Dai W, Kloner RA, Pahlevan NM. A Physics-Based Machine Learning Approach for Instantaneous Classification of Myocardial Infarct Size. Circulation. 2021;144(Suppl_1):A12098–A.

45. Pahlevan NM, Alavi R, Ramos M, Hindoyan A, Matthews RV. An Artificial Intelligence Derived Method for Instantaneous Detection of Elevated Left Ventricular End Diastolic Pressure. Circulation. 2020;142(Suppl_3):A16334–A.

46. Alonso A, Aparicio FHJ, Benjamin EJ, Bittencourt MS, Callaway CW, Carson FAP, et al. Heart Disease and Stroke Statistics—2021 Update. Circulation. 2021;2021(143):e00–e.

47. Rafieianzab D, Abazari MA, Soltani M, Alimohammadi M. The effect of coarctation degrees on wall shear stress indices. Scientific Reports. 2021;11(1):1–13.

